# Composition and Functional Properties of Hemp Seed Protein Isolates from Various Hemp Cultivars

**DOI:** 10.1101/2022.06.01.494437

**Authors:** Martin Liu, Jacob A. Toth, Mackenzie Childs, Lawrence B. Smart, Alireza Abbaspourrad

## Abstract

Hemp seed protein isolates (HPI) were extracted from the seeds of seven commercial hemp cultivars, a Cornell breeding line, and a commercial hemp heart product. The composition and functional properties of the resulting HPI were investigated. HPI were of high protein purity >96% and contained various ratios of the major protein classes edestin, vicilin, and albumin, depending on the source. Protein solubility varied across HPI, and there was a positive correlation between greater levels of vicilin and albumin and improved solubility. The isoelectric points of HPI ranged from 5.50 to 5.94 but did not show significant effects associated with major protein class ratio. Significant differences in HPI foam capacity (52.9–84.9%), foam stability (68.1–89.4%), water holding capacity (0.83–1.05 g water/g protein isolate), and oil holding capacity (1.28–1.81 g oil/g protein isolate) were observed. In general, the emulsions generated from HPI performed poorly in terms of emulsifying activity, stability, and emulsion particle size. The ratio of edestin, vicilin, and albumin was found to play a major role in HPI functionality, suggesting that certain hemp cultivars may be better suited for generating protein ingredients that can functionalize plant-based foods.

## 1. Introduction

Plant protein-based foods have gained in popularity and market share compared to animal protein-based foods due to their environmental benefits, positive health implications, and improved product development (Willett et al., 2019). Plant protein ingredients are commonly sourced from soy, pea, wheat, and beans, and confer macronutrients and functional properties to their final food products (McClements & Grossmann, 2021). These proteins hold onto water and oil and stabilize colloidal systems including emulsions, foams, and gels (Loveday, 2020). However, common allergens associated with these plant sources hinders consumer adoption (Hertzler, Lieblein-Boff, Weiler, & Allgeier, 2020), and the lack of thoroughly researched protein ingredients limits product development for more consumer-accepted plant-protein foods.

Hemp seed is a promising, yet understudied, source of plant protein ingredients that has only been utilized in a few alternative dairy food products. Hemp is defined as *Cannabis sativa* which produces less than 0.3% of the psychoactive compound tetrahydrocannabinol (Congressional Research Service, 2019). Hemp is capable of generating large seed yields with high protein concentrations (House, Neufeld, & Leson, 2010) and without gluten and legume allergens. However, hemp seed protein suffers from poor solubility at neutral pH, hindering its utility as a functional food ingredient (Wang & Xiong, 2019). Downstream processing techniques to improve hemp seed protein functionality have been explored, including dehulling (Shen et al., 2020), salt extraction (Hadnađev et al., 2018), and pH shift (Wang, Jin, & Xiong, 2018), but these methods may have an upper limit to their improvement on protein functionality. In order to become competitive with other plant protein ingredients, hemp seed may benefit from genetic improvements to resolve these functional setbacks.

The goal of this study is to draw connections between the composition and functional properties of hemp seed protein isolate (HPI), which is composed of three protein groups: edestin, vicilin, and albumin (Wang & Xiong, 2019). Variations in protein group ratio have previously been linked to changes in protein functionality across cultivars of pea (Cui et al., 2020; Lam, Warkentin, Tyler, & Nickerson, 2017; Stone, Avarmenko, Warkentin, & Nickerson, 2015), chickpea (Chang et al., 2022), and wheat (Barak, Mudgil, & Khatkar, 2015), but not yet for hemp. Herein, HPI protein group relative composition and functional properties are investigated, and relationships between edestin, vicilin, and albumin ratios and protein functionality are examined by principal component analysis. Differences in protein functionality underscore the importance of hemp seed sourcing and suggest that there is potential to enable hemp seed protein utilization in plant-based foods through breeding.

## 2. Materials & Methods

### 2.1 Materials

Hemp seed from the commercial cultivars ‘Anka’ (A), ‘Bialobrzeskie’ (B), ‘Hlukhivs’ki 51’ (H), ‘Picolo’ (P), and ‘Yuma Crossbow’ (Y) and Cornell hemp breeding line GVA-H-20-1179 (G) was harvested from Cornell hemp research field trials grown in 2020. Hemp seed of commercial cultivars ‘La Crème’ (L) and ‘Magic Bullet 5’ (M) was donated by Point3Farma (Center, CO, USA). Commercial hemp hearts (C) were purchased from Manitoba Harvest (Winnipeg, Canada). Raw hemp seeds were dehulled with a Forsberg (Thief River Falls, MN, USA) Model 7-F impact huller and expeller pressed with a Kern Kraft (Reut, Germany) KK20 F Universal unit. The resulting hemp seed press cakes were frozen at -20 °C until further use.

Hexanes, disodium phosphate, monosodium phosphate, and acetic acid were purchased from Fisher Chemical (Waltham, MA, USA). Hydrochloric acid, sodium chloride, citric acid, sodium citrate, sodium dodecyl sulfate, and methanol were purchased from Sigma-Aldrich (St. Louis, MO, USA). Tris, glycine, and Coomassie brilliant blue R-250 were purchased from Bio-Rad (Hercules, CA, USA). Sodium hydroxide was purchased from Bio Basic (Markham, Canada) and corn oil from a local grocery store. All chemicals were used as received.

### 2.2 Preparation of protein isolates

Hemp seed press cakes were defatted in 20 % (w/v) hexanes under mechanical stirring at room temperature for 2 h. The resulting slurry was vacuum filtered, rinsed with additional hexane until the eluent ran clear, and dried in an oven at 40 °C overnight. Defatted hemp seed material was ground to a fine powder with a coffee mill.

Alkaline extraction and isoelectric precipitation of HPI was performed based on previous methods (Tang, Ten, Wang, & Yang, 2006) with recommended changes (Potin, Lubbers, Husson, & Saurel, 2019) and some additional minor modifications. Defatted hemp seed powder was suspended in deionized water at 10% (w/v), and the pH of the solution was adjusted to 10.0 with 1 M NaOH. The suspension was stirred at 35 °C for 1 h, and at the 20 and 40 min marks, the pH was re-adjusted to 10.0. After centrifugation at 8000×*g* for 30 min at room temperature, the supernatant was adjusted to a pH of 5.0 with 1 M HCl and centrifuged at 8000×*g* for 10 min at 4 °C. Following decantation, the pellet was resuspended in deionized water, frozen at -20 °C, and lyophilized at -52 °C.

### 2.3 Protein quantification

Protein concentration was determined by AOAC method 991.20 for total nitrogen content (AOAC, 2016). The total nitrogen content was multiplied by 6.25 in order to calculate the total (N x 6.25) protein content in sample materials. Sample size was adjusted to achieve about 0.3 g protein per analysis for the material in this study. Experiments were performed in duplicate.

### 2.4 Gel electrophoresis

Sodium dodecyl sulfate – polyacrylamide gel electrophoresis (SDS-PAGE) gels were prepared using a TGX Stain-Free FastCaste Acrylamide Kit, 12 %, from Bio-Rad (Hercules, CA, USA). The running buffer contained 2.5 mM Tris, 19.2 mM glycine, and 0.1 % SDS (w/v). Protein isolates were solubilized at 0.15 % (w/v) in 10 mM Tris buffer (pH 10.0) and mixed at a 1:1 ratio with 2x Laemmli sample buffer (62.5 mM Tris-HCl at pH 6.8 with 2 % SDS, 25 % glycerol, and 0.01 % bromophenol blue (w/v)) and loaded onto the gel. Bio-Rad Precision Plus Protein Dual Color Standards (Hercules, CA) molecular weight ladders were run alongside the protein samples. Electrophoresis was performed at 200 V for 45 min, and the resulting gel was stained (50 % methanol, 10 % acetic acid (v/v), 0.15 % Coomassie Brilliant R-250 (w/v)) for 30 min. The gel was destained overnight with three exchanges of the destaining buffer (20 % methanol, 10 % acetic acid (v/v)).

Destained gels were imaged on a Bio-Rad ChemiDoc Imaging System (Hercules, CA, USA), and images were analyzed using Bio-Rad ImageLab software (version 6.1). Quantification of band saturation was conducted similarly to that of Lam et al. (2017), but with polypeptides selected to represent the hemp seed proteins vicilin (∼47,000 Da), edestin (∼33,000 Da [acidic subunit]), and albumin (∼10,000 Da). Pixel saturation intensity from all three bands was summed for each well on the SDS-PAGE gel, and ratios of each polypeptide was calculated from this total.

### 2.5 Zeta-potential and isoelectric point analysis

Zeta-potential experiments were conducted using a Malvern Zetasizer Nano ZS90 (Malvern, UK) with 0.05 % (w/v) dispersions of HPI in water at room temperature. The pH of the sample was adjusted to 10.0 with 1 M NaOH, and the surface charge was measured. A refractive index of 1.330 for water was input into the software, and a minimum of 10 measurements were made for each sample. The pH was then adjusted incrementally (9.0, 8.0, 7.0, … 2.0) with 1 M HCl, and zeta-potentials were measured at each point. Each cultivar was measured for zeta-potential profiles in duplicate.

Isoelectric points were calculated by linear approximation of the lowest positive and negative value measurement for each zeta-potential profile. A line segment between the two zeta-potential data points was drawn, and the x-intercept was calculated for each cultivar in duplicate.

### 2.6 Protein solubility

Hemp seed protein isolates were dispersed at 1 % (w/v) in 50 mM citrate (pH 3.0, 4.0, 5.0, 6.0) and phosphate (pH 7.0) buffers overnight at room temperature. Following centrifugation at 10,000×*g* for 20 min, the supernatant was analyzed for protein concentration by the Lowry method (Lowry, Rosebrough, Farr, & Randall, 1951). Solubility is expressed as a percentage of protein in the supernatant compared to the original amount of protein isolate dispersed.

### 2.7 Foaming properties

Foams were prepared with 1 % (w/v) dispersions of HPI in 5 mL of phosphate buffer (pH 7.0) overnight at room temperature. Using an IKA T25 Ultra Turrax homogenizer equipped with an S25N-8G dispersing element (Staufen, Germany), samples were foamed at the air-water interface at 20,000 rpm for 5 min. Foaming capacity is calculated as a percentage of the foam volume generated from the initial dispersion volume. Foam stability is calculated as a percentage of the remaining foam volume after 30 min compared to the initial foam volume.

### 2.8 Emulsifying properties

Emulsions were prepared with 1% (w/v) dispersions of HPI in 4.5 mL of phosphate buffer (pH 7.0) overnight at room temperature. After adding 0.5 mL of corn oil, the sample was homogenized with an IKA T25 Ultra Turrax homogenizer equipped with an S25N-8G dispersing element (Staufen, Germany) at the oil-water interface at 20,000 rpm for 5 min.

Turbidimetric methods for evaluating emulsions were based on the methods by Pearce & Kinsella with modifications (Pearce & Kinsella, 1978). Thirty minutes after homogenization, emulsions were sampled below the cream layer, diluted (1:100, v/v) in 0.1 % (w/v) SDS solution, and vortexed for 5 s. The absorbance of the diluted emulsion was recorded at 500 nm with a Shimadzu UV-2600 spectrophotometer (Kyoto, Japan). The reference absorbance with SDS solution was subtracted to correct for sample absorbance. Emulsifying activity index (EAI) and emulsion stability index (ESI) were calculated by the following equations:

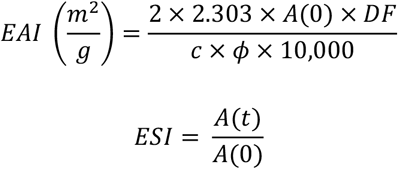

where A(0) is the absorbance 30 min after emulsion formation, A(t) is the absorbance after 48 h, DF is the dilution factor (100), c is the protein concentration (0.01), and φ is the oil volume fraction of the emulsion (0.05).

Emulsion droplet size after 48 h of storage was determined using a Malvern Zetasizer Nano ZS. Emulsions were diluted (1:10, v/v) in the sample buffer and inverted thrice to mix. The refractive indexes of corn oil and water were 1.473 and 1.330, respectively. Average emulsion particle size (Z-Ave) was measured in triplicate.

### 2.9 Water and oil holding capacities

Water holding capacity (WHC) and oil holding capacity (OHC) were determined by suspending 500 mg of HPI in 5.0 g of deionized water or corn oil in a centrifuge tube, vortexing for 1 min, and letting it stand for 30 min. The samples were centrifuged at 7000×*g* for 25 min at room temperature, and the supernatant was drained by inverting the tube for 15 min. The remaining pellet was weighed, and WHC/OHC were calculated by dividing the additional weight of the pellet by the original protein isolate mass.

### 2.10 Statistical analysis

Analysis of variance (ANOVA) was performed on JMP Pro 16 statistical software (Cary, NC), and letters denote statistically significant differences by means of Tukey’s test (p < 0.05). Principal component analysis (PCA) was performed in R version 3.5.1 (R Core Team 2018), and correlation data was visualized using the corrplot library (Wei & Simko, 2021).

## 3. Results and Discussion

### 3.1 Protein composition in raw hemp seeds and isolates

High protein concentration is an important factor in assessing the viability of processing hemp seed as a source of plant protein ingredients. Among the eight samples of raw hemp seed analyzed, protein concentration spanned a range of 24.20–27.68 % (Table 1). Hemp seed contained less protein than soybeans (33.0–39.0 %) (Assefa et al., 2019) and pea (28–30 %) (Lu, He, Zhang, & Bing, 2020) but more than chickpea (19.20–22.50 %) (Deep Singh, Wani, Kaur, & Sogi, 2008) and wheat (10.1–15.7 %) (Giunta, Pruneddu, & Motzo, 2019). Following alkaline extraction-isoelectric precipitation, the generated HPIs exhibited high protein purity > 96 %. Overestimation of protein concentration (> 100 %) was noted in several HPI samples and may be attributed to an inaccurate Kjeldahl conversion factor. Nitrogen to protein ratios depend on the protein source, and while 6.25 is the standard factor used across most foodstuffs, plant proteins may require factors with much lower values (Mariotti, Tomé, & Mirand, 2008). Oilseeds are recommended 5.30 as a stand-in factor (Jones, 1941), but further studies to estimate a specific conversion factor for hemp seed are needed.

**Table 1.** Protein concentration of raw seeds and protein isolates from various sources of hemp seeds.

| Source | Abbreviation | Protein concentration (%) |  |
| --- | --- | --- | --- |
|  |  | Raw seeds | Protein isolate |
| Piccolo | P | 27.68 ± 0.26 a | 96.65 ± 0.15 d |
| Magic Bullet 5 | M | 27.60 ± 0.41 a | 100.64 ± 0.48 a |
| La Crème | L | 27.38 ± 0.28 ab | 99.22 ± 0.17 b |
| Hlukhivs'ki 51 | H | 26.58 ± 0.15 bc | 97.82 ± 0.41 c |
| Bialobrzeskie | B | 25.85 ± 0.12 cd | 97.19 ± 0.11 cd |
| Yuma Crossbow | Y | 25.29 ± 0.12 de | 96.66 ± 0.40 cd |
| Anka | A | 24.68 ± 0.25 ef | 100.31 ± 0.30 ab |
| GVA-H-20-1179 | G | 24.20 ± 0.28 f | 96.38 ± 0.25 d |
| Commercial hemp hearts | C | - | 101.02 ± 0.04 a |

Protein concentration values represent the mean ± one standard deviation (n = 2), and different letters within a column represent statistically significant differences as calculated by Tukey's HSD (p < 0.05).

Hemp seed protein isolates displayed similar SDS-PAGE profiles containing the three major hemp seed proteins (edestin, vicilin, and albumin) with a few differences in minor polypeptide bands (Figure 1A). The acidic and basic subunits of edestin resolved at 33 kDa and 18-22 kDa, vicilin subunits at 47 kDa, and albumin polypeptides at 10 kDa (Wang & Xiong, 2019). A minor polypeptide band at 55 kDa was missing in sample G, and another at 27 kDa was missing in samples C and M. These variations in polypeptides may be due to the genetic diversity in hemp seed proteins, as multiple genes that code for edestin and albumin have previously been identified (Docimo, Caruso, Ponzoni, Mattana, & Galasso, 2014; Ponzoni, Brambilla, & Galasso, 2018).

**Figure 1.**
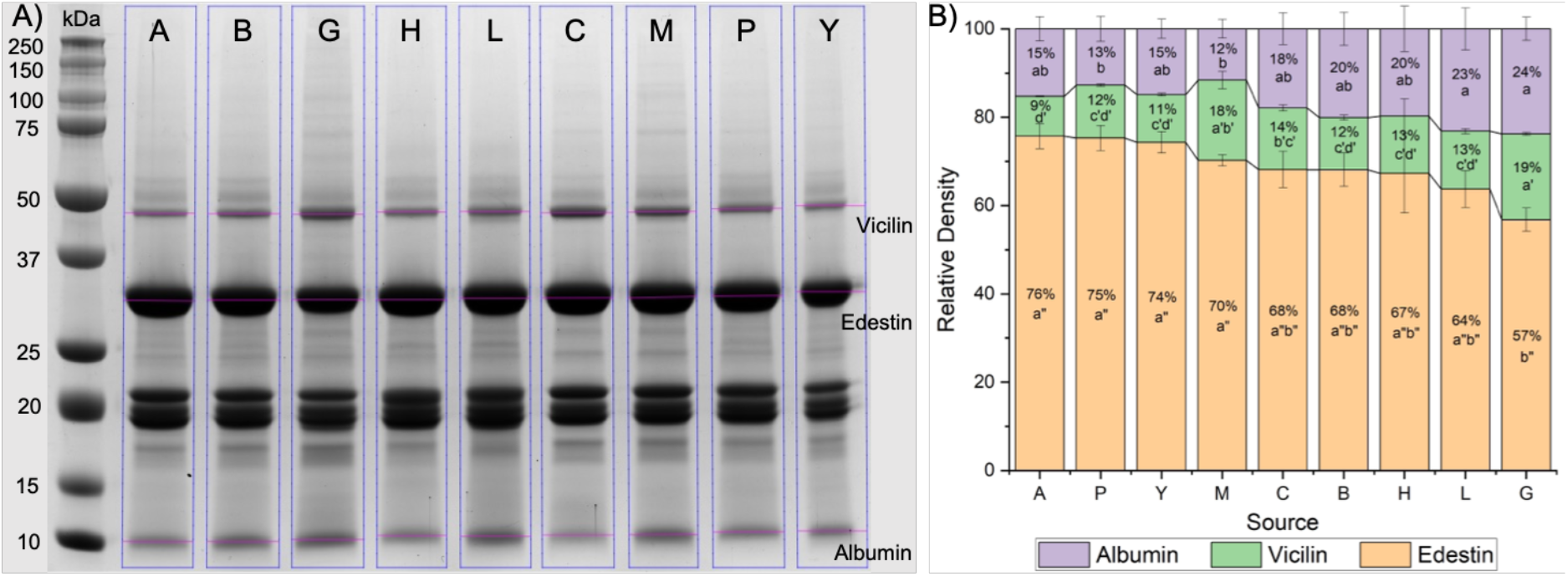
(A) Representative reducing SDS-PAGE gel with HPI. Band saturation as a proxy for protein concentration was quantified on ImageLab and graphed as (B) relative densities of edestin, vicilin, and albumin. Three gels were prepared and quantified independently, error bars represent one standard deviation from the mean, and different letters with the same prime notation represent statistically significant differences as calculated by Tukey’s HSD (p < 0.05).

Pixel saturation in the SDS-PAGE gels was used as a proxy for calculating the relative densities of edestin (33 kDa), vicilin (47 kDa), and albumin (10 kDa). Three gels were analyzed, and their mean values are reported (Figure 1B). Due to the polydisperse nature of the bands that represent the basic subunits of edestin, only the acidic subunits of edestin at 33 kDa were measured. In each protein isolate, the majority of protein was from edestin, a finding that has been reported in the literature (Sun, Sun, Li, Wu, & Wang, 2021). The relative proportion of edestin varied from 76 % in A down to 57 % in G. Albumin comprised the second greatest protein fraction, between 12-24 % of relative protein density, while vicilin contributed the least amount of protein, between 9-19 %. The sum of vicilin and albumin was inversely proportional to the relative density of edestin in HPI, suggesting that different sources of hemp seed that undergo the same protein extraction process resulted in different ratios of proteins in the isolate.

### 3.2 Protein solubility and isoelectric points

Solubility is an important characteristic that can profoundly influence the functionality of proteins for stabilizing colloidal systems. Many foods and beverages are formulated in neutral to acidic conditions, often with buffering systems, so the solubility of HPI at buffered pH between 3.0 and 7.0 was investigated (Figure 2A). In general, HPI was most soluble at pH 3.0 and showed similar solubilities between pH 4.0 to 7.0. Within a given pH, statistically significant differences in protein solubility can be observed across cultivars (Table S1). At pH 7.0, samples G and L displayed greater solubility, 17.50 and 14.50 %, respectively, while the other HPI had lower solubility.

**Figure 2.**
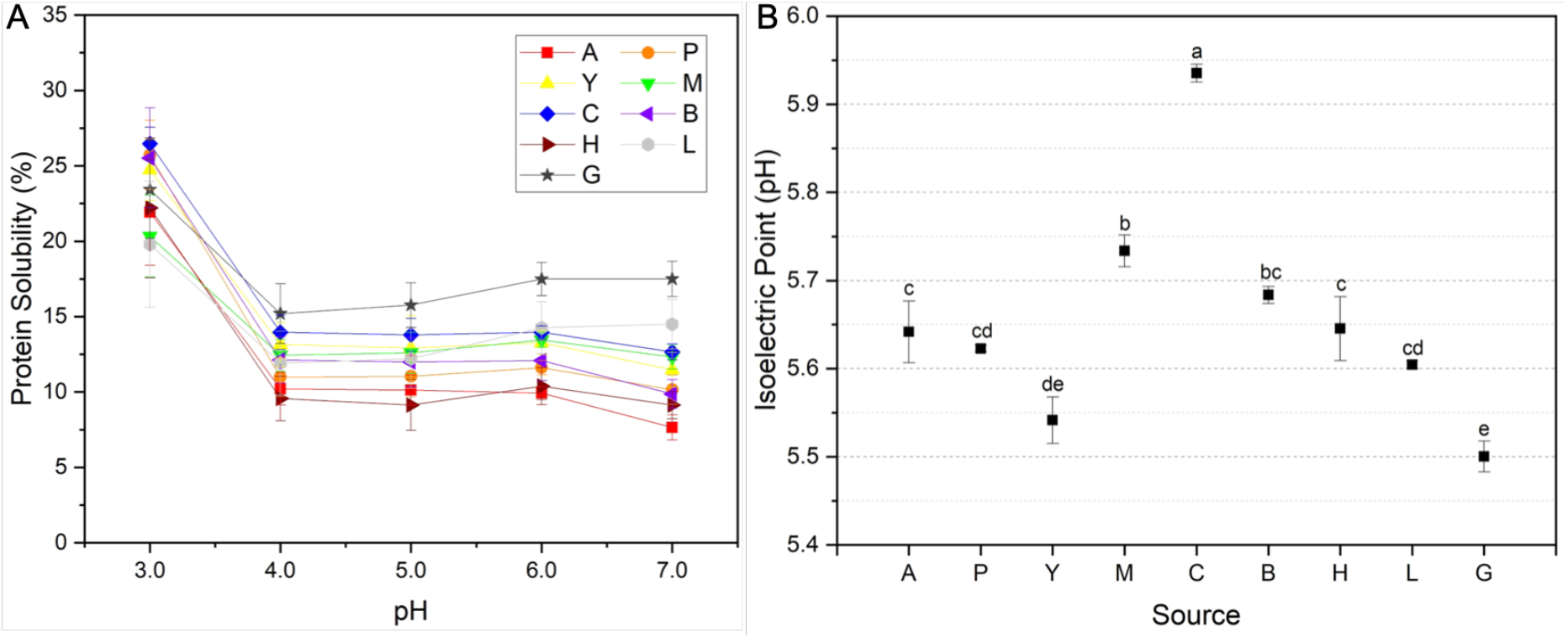
(A) Protein solubility of HPI solubilized in pH 3.0-7.0 buffers. Detailed values and statistical analysis can be found in Table S1. Experiments were performed in triplicate, and error bars represent one standard deviation from the mean. (B) Isoelectric points for HPI calculated by linear approximations of zeta-potential profiles displayed in Figure S1. Experiments were conducted in duplicate, error bars represent one standard deviation from the mean, and different letters represent statistically significant differences as calculated by Tukey’s HSD (p < 0.05).

The solubility curves reported in Figure 2A differ from those that have previously been reported. Compared to data from Shen et al., our HPI were lower in solubility from pH 3.0-5.0, higher in solubility at pH 6.0, and approximately the same at pH 7.0 (Shen et al., 2020). Rather than the typical “u-shape” that is typical of plant proteins, the solubility of our samples plateaued as the environmental pH approached 7.0. The difference in findings may be due to our buffered methodology for measuring protein solubility. Buffers play an integral role in stabilizing protein formulations, and various buffering systems can have stabilizing or destabilizing effects on proteins in solution. These effects are well known in pharmaceutical settings but understudied in the context of food. For example, protein solubility of most HPI samples reduced when pH increased from 6.0 to 7.0. This may seem counterintuitive due to a hypothesized increased surface charge as the environmental pH deviates from the isoelectric point of hemp seed protein ∼ pH 5.0 (Hadnađev et al., 2018; Tang et al., 2006). Sodium citrate buffer was used to measure protein solubility up to pH 6.0, but since the buffering system does not function above pH 6.2, sodium phosphate buffer was used to measure protein solubility at pH 7.0. Citrate, with its multiple carboxylate groups, can stabilize proteins by ligand binding and improving conformational stability (Zbacnik et al., 2017). These matrix effects may explain the lack of a minimum solubility point that is characteristic of a typical “u-shape” solubility curve, but further studies would be needed to investigate the mechanisms of buffer-protein interactions.

The surface charge characteristics of HPI followed similar patterns across all samples measured (Figure S1). Zeta-potential, a measurement of surface charge, is influenced by the pH of the dissolving solution and structural characteristics of the protein itself. In general, the lowest zeta-potential values were observed at a pH of 10.0 (∼ -20 to -34 mV), and the greatest values were observed at a pH of 3.0 or 4.0 (∼+13 to +22 mV). At pH 2.0, zeta-potentials in every sample were slightly lower than maximum values, a phenomenon that has previously been reported in hemp seed protein (Shen et al., 2020) and may be a result of acid hydrolysis from glutamine and/or asparagine residues into glutamic acid and/or aspartic acid (Ladjal-Ettoumi, Boudries, Chibane, & Romero, 2016).

The isoelectric point of a protein occurs when the environmental pH causes the surface charge of the protein to equal to net zero, typically resulting in protein aggregation and diminished solubility. Isoelectric points were estimated by linear approximation of the two points closest to zero for each zeta-potential profile (Figure 2B). Most HPI exhibited isoelectric points between pH 5.6 and 5.7, but the range spanned 5.50 in G up to 5.94 in C. This is in contrast to previously reported isoelectric points of pH 5.0 typically used in protein extraction by isoelectric precipitation (Hadnađev et al., 2018; Tang et al., 2006), but in general agreement with the isoelectric point of 5.8 reported by Shen et al. (Shen et al., 2020).

### 3.3 Functional properties

#### 3.3.1 Foaming properties

The ability of HPI to generate and stabilize foams at pH 7.0 varied depending on the source of hemp seed (Figure 3). Foam capacity ranged from 52.9 % for sample P up to 80.4 % and 84.9 % for samples G and L, respectively. The resulting foams were generally stable, retaining between 68.1–89.4 % of the original foam volume after 30 min. There does not appear to be any correlation between foam capacity and foam stability. Variability in foam capacity and stability between HPI indicates that certain sources may be more suitable than others for food foaming applications.

**Figure 3.**
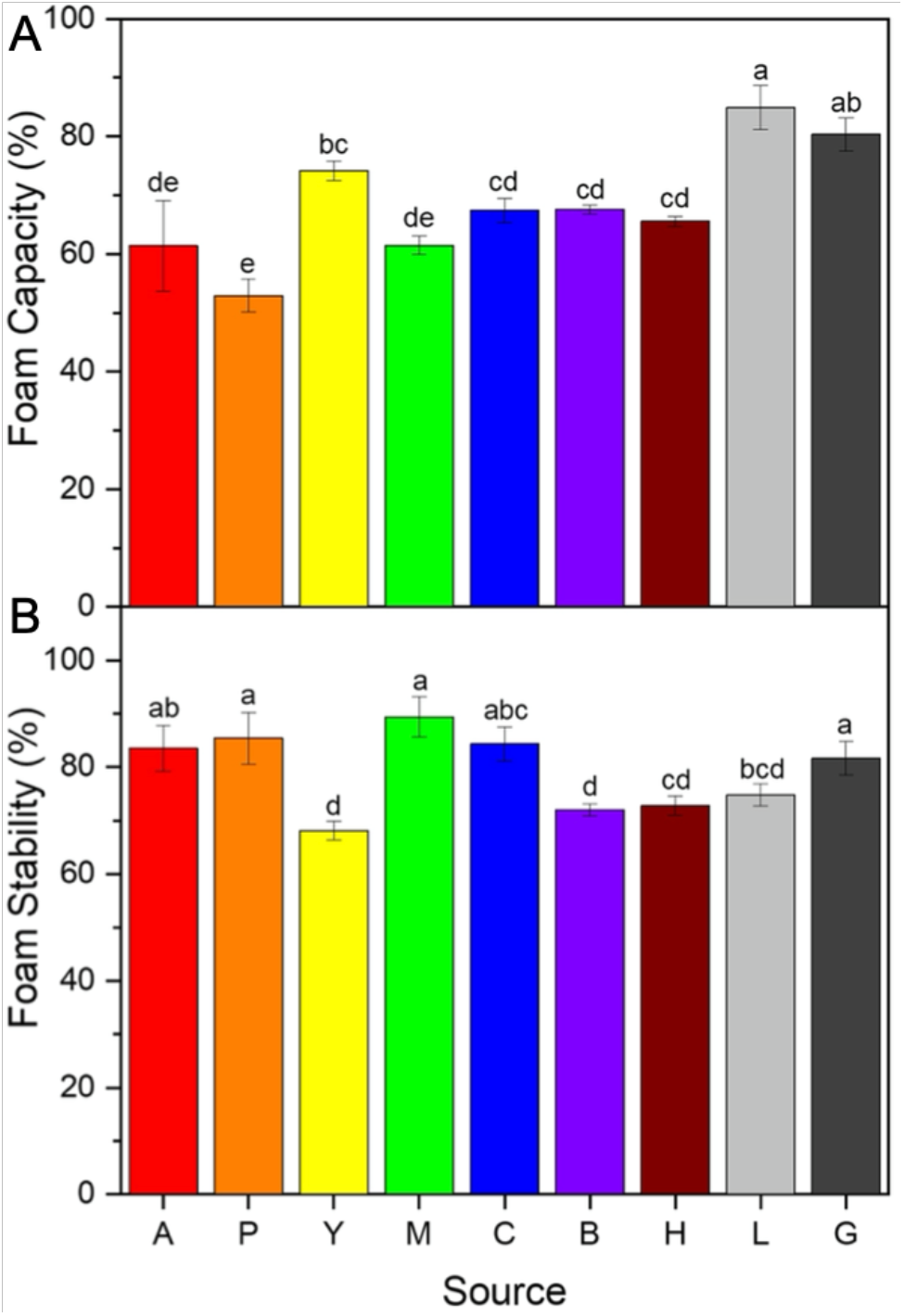
Foaming properties of HPI at pH 7.0 in terms of (A) foam capacity and (B) foam stability after 30 min left undisturbed. Experiments were conducted in triplicate, error bars represent one standard deviation from the mean, and different letters represent statistically significant differences as calculated by Tukey’s HSD (p < 0.05).

The emulsifying properties of HPI were uniform across all samples with a few exceptions (Figure 4). EAI for most protein isolates was between 1.1 and 1.7 m^2^ / g with no statistically significant differences among samples, other than sample G, which exhibited twice the emulsifying activity at 3.5 m^2^ / g. After 48 h of storage, ESI of the HPI emulsions was measured, ranging from 38–73 % but large standard deviations negated any significant differences. The mean diameter of the emulsion droplets was also measured between 40–90 µm, but large standard deviations also prevented the identification of significant differences among samples. Overall, low EAI values and large standard deviations across all emulsion metrics indicate that HPI was not a good emulsifying agent, though the elevated EAI value for sample G is noteworthy and may hint at potential room for improvement.

**Figure 4.**
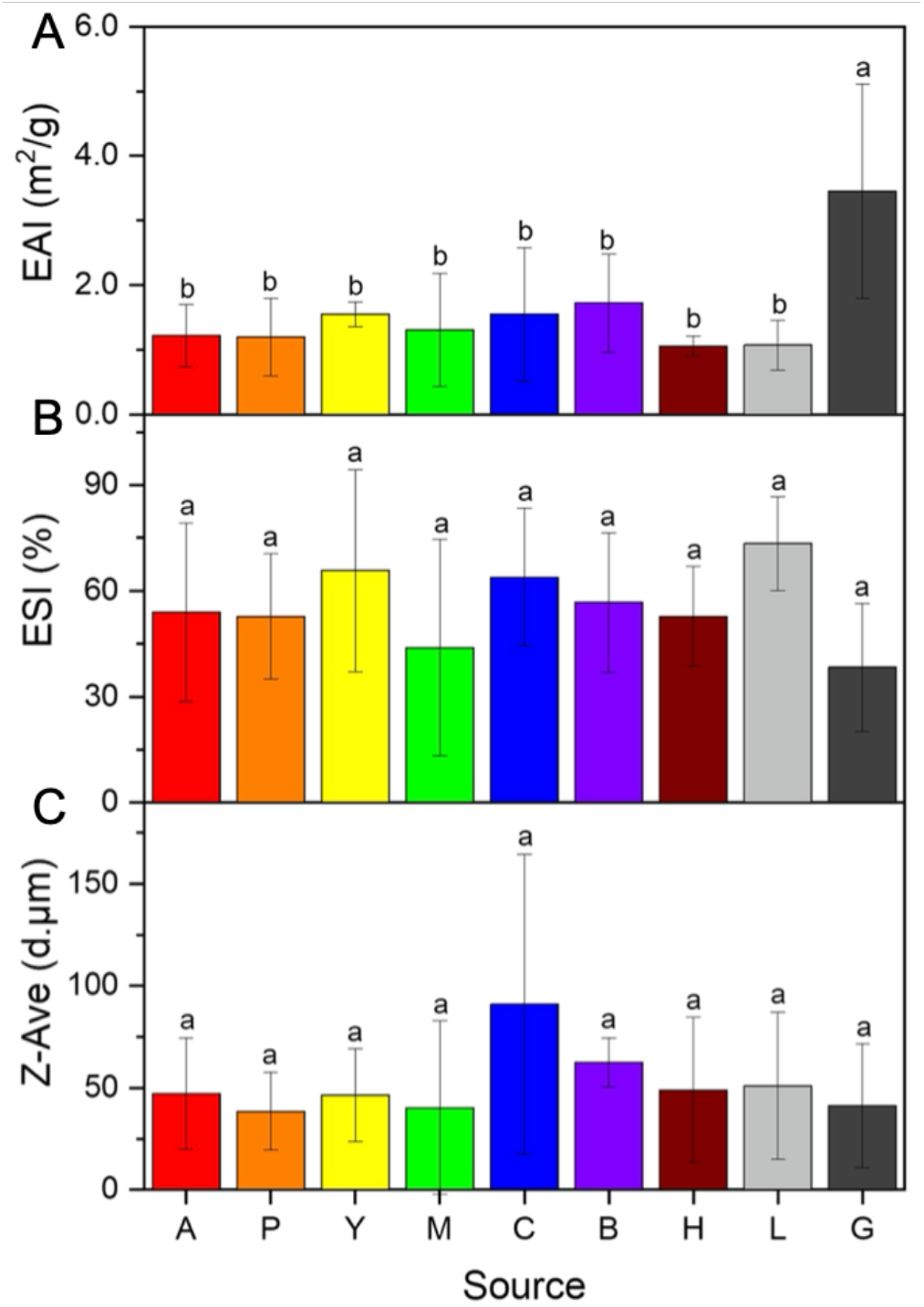
Emulsifying properties of HPI at pH 7.0 in terms of (A) emulsifying activity index (EAI), (B) emulsifying stability index (ESI), and (C) average emulsion particle size (Z-Ave). Experiments were conducted in six replicates, error bars represent one standard deviation from the mean, and different letters represent statistically significant differences as calculated by Tukey’s HSD (p < 0.05).

The fluid holding capacity of HPI varied depending on the source of hemp seed (Figure 5). WHC for the protein isolates ranged between 0.83 – 1.05 g water / g isolate, with sample A exhibiting the greatest capacity to retain water and sample L the lowest. Oil holding capacity (OHC) spanned an even wider range from 1.28 – 1.81 g corn oil / g isolate, with sample P displaying 41 % greater OHC over sample M. Compared to pea protein isolate, HPI exhibited a poorer WHC but better OHC (Stone et al., 2015). This comparison, along with a large observed difference in OHC among HPI, indicates that certain sources of hemp seed can produce protein isolates with good oil binding abilities.

**Figure 5.**
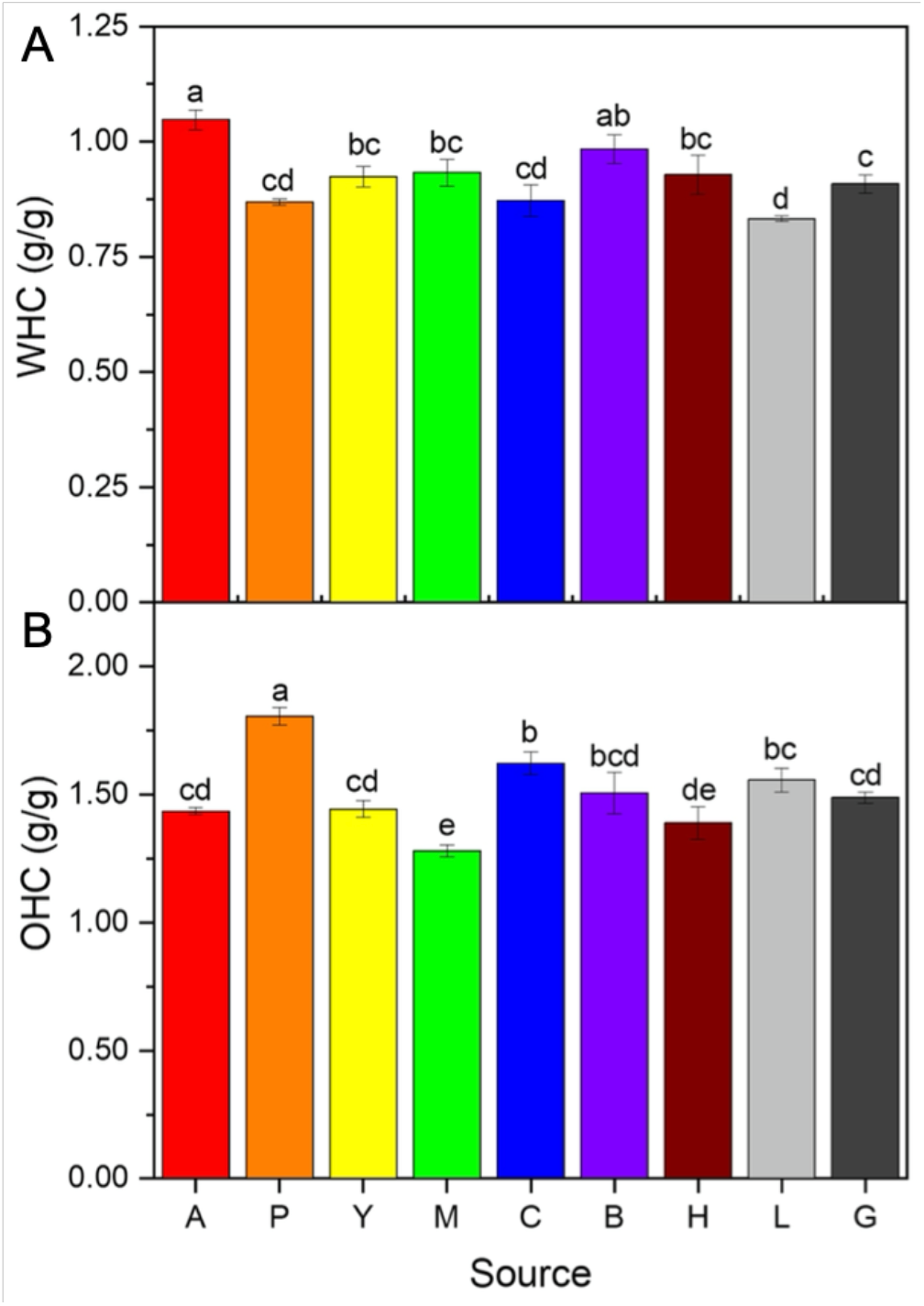
(A) Water holding capacity (WHC) and (B) oil holding capacity (OHC) for HPI. Experiments were conducted in triplicate, error bars represent one standard deviation from the mean, and different letters represent statistically significant differences as calculated by Tukey’s HSD (p < 0.05).

### 3.4 Principal component analysis

Principal component analysis was used to visualize the patterns of covariation between hemp seed sources and the properties of HPI that were evaluated in this study (Figure 6A). The first principal component (PC1) explained 41.8 % of the variance and can be generalized as associations with albumin and vicilin versus associations with edestin. Sample G was clearly separated from other hemp seed sources by PC1 alone, reflecting a distinct difference in protein composition. The second principal component (PC2) represents 17.4 % of data variance and further distinguished HPI based on functional properties.

**Figure 6.**
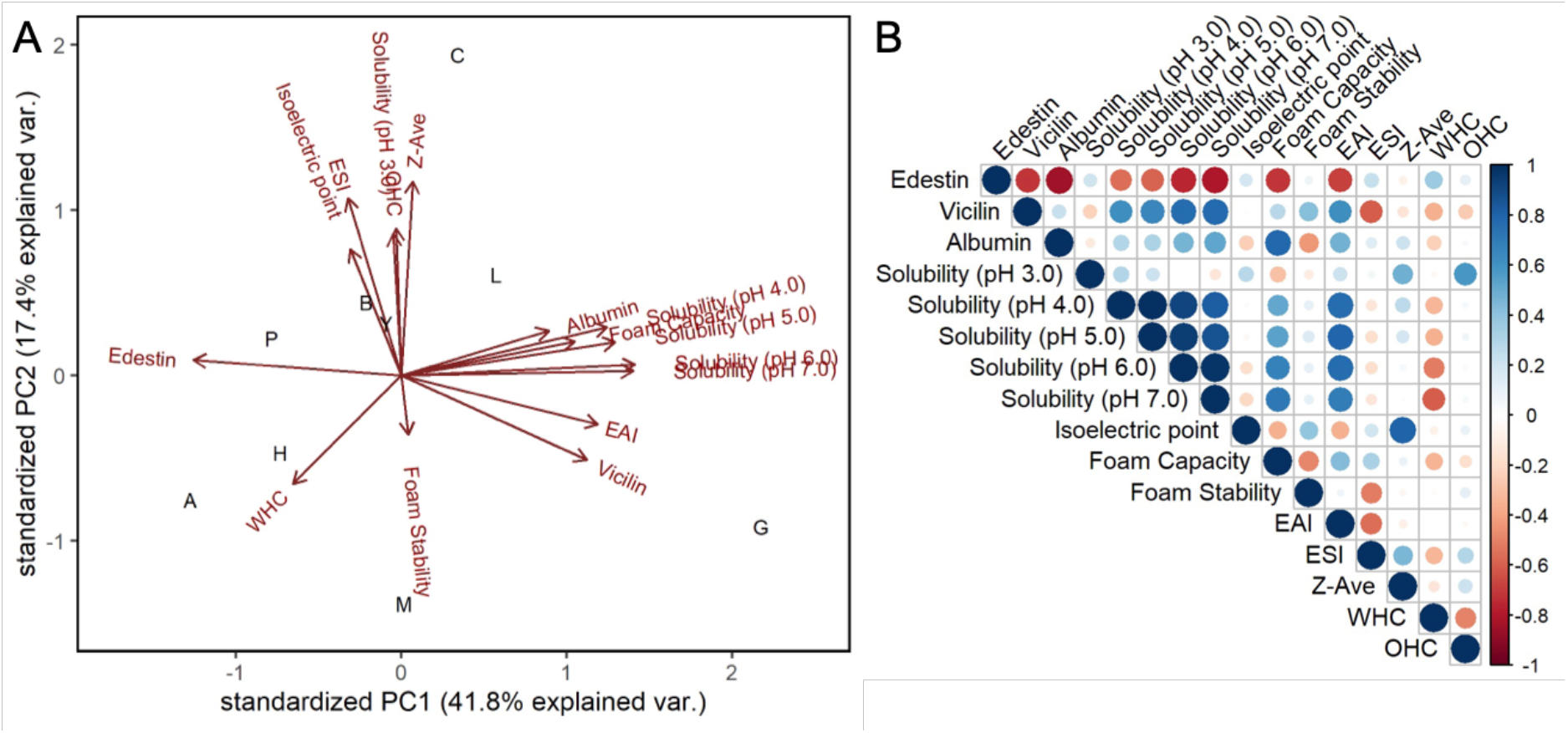
(A) Combined score and loading plots for principal component analysis (PCA) conducted on the HPI and the correlations between parameters measured in this study. (B) Correlation (Pearson’s R) between protein composition and functional properties of HPI. Positive correlation is depicted by blue circles, negative correlation is depicted by red circles, and no correlation is depicted with empty squares. EAI: emulsifying activity index; ESI: emulsifying stability index; Z-Ave: average emulsion particle size; WHC: water holding capacity; OHC: oil holding capacity.

### 3.5 Relationships between protein composition and functionality

Significant correlations between HPI composition and functional properties were identified (Figure 6B). Protein composition had a direct and significant impact on functional properties of protein isolates, so variation in protein relative densities across hemp seed sources resulted in differences in functional properties. This composition-function relationship has been demonstrated in cultivars of pea (Cui et al., 2020; Lam et al., 2017; Stone et al., 2015), chickpea (Chang et al., 2022), and wheat (Barak et al., 2015).

The proportion of edestin was strongly and inversely related to the proportions of vicilin and albumin, such that their zero-sum relationship can also explain variations in protein functional properties. The proportion of edestin was also inversely correlated with protein solubility at pH 4.0–7.0. Our data are consistent with previous findings that hemp seed protein globulins, the protein fraction that contains edestin, exhibits poor solubility due to its isoelectric point at pH 5.0, higher levels of aromatic and hydrophobic amino acids, and extreme protein tertiary structure changes across pH values (Malomo & Aluko, 2015). Foam capacity and EAI were also negatively associated with high edestin concentrations, which may be partially explained by the low solubility of edestin-rich protein isolates at pH 7.0.

The proportions of vicilin and albumin were positively correlated with improved solubility at pH 4.0–7.0, though this relationship was stronger for vicilin than for albumin. Both protein fractions were also associated with improved foam capacity, but albumin contributed a larger role possibly due to its smaller size and its interactions with the aqueous phase that aids in foam generation. However, albumin conferred a decrease in foam stability because of weaker films formed at the air-water interface, which led to bubble coalescence and foam destabilization (Malomo & Aluko, 2015). Vicilin and albumin are associated with improved EAI via increased protein solubility, since the formation of emulsions necessitates the solubilization of oil-water interface stabilizing moieties. Vicilin appears to be negatively correlated with ESI, but our data did not show significant differences in ESI values across HPI samples, so this relationship cannot be similarly assessed.

Fluid holding capacity was not strongly associated with protein composition, but rather, several other functional properties. WHC appeared to be negatively correlated with solubility at pH 4.0–7.0, but this may be due to methodology. Protein isolates that are generally more soluble will lose more dissolved mass when water is decanted from the sample. Changes to the method, such as reducing the amount of water mixed with the protein isolate, could reduce this error. WHC seemed to have an inverse relationship with OHC, indicating that different sources of HPI may offer improved holding capacity for water or oil, but not typically both.

## 4. Conclusion

The composition and functional properties of protein isolates extracted from different sources of hemp seed were investigated. Although the raw hemp seeds expressed similar levels of protein, the HPI extracted by alkaline extraction-isoelectric precipitation contained varying proportions of edestin, vicilin, and albumin. Isoelectric points for the HPI were higher than previously reported in literature. Protein solubility of the isolates was also different than previous reports, possibly due to matrix effects from the buffering systems used, and HPI from the Cornell breeding line consistently exhibited improved solubility over the other samples due to its greater ratio of vicilin and albumin to edestin. Foam capacity was also higher in samples with a greater proportion of vicilin and albumin to edestin, but foam stability was negatively impacted by albumin concentration. EAI for HPI from the Cornell breeding line was markedly greater than all others, but overall low EAI values and large standard deviations across ESI and Z-Ave indicate poor emulsifying ability for HPI. Significant differences in WHC and OHC across the HPI demonstrate the differences in fluid holding capacity due to hemp seed source.

Moderate foam capacity and stability and a high OHC indicate the possible utilization of HPI for targeted functional applications in plant-based foods. The variable ratios of edestin, vicilin, and albumin across sources of HPI and their direct impact on protein functionality, suggest breeding and selection for improved ratio of vicilin and albumin to edestin may result in hemp cultivars with improved protein functional properties.

## Acknowledgements

The authors would like to acknowledge the Barbano Research Laboratory, including Dr. David Barbano, Michelle Bilotta, and Chassidy Coon, in the Department of Food Science at Cornell University for conducting Kjeldahl analysis. We are grateful to Uniseeds, Hemp Genetics International, International Hemp, Fiacre Enterprises, Point3 Farma, and Harvey Farms for providing hemp seed for trials. This work was partially funded by the New York State Department of Agriculture and Markets through grants from Empire State Development Corporation (AC477, AC483, and #132,997).

## Supplementary Information

**Figure S1.**
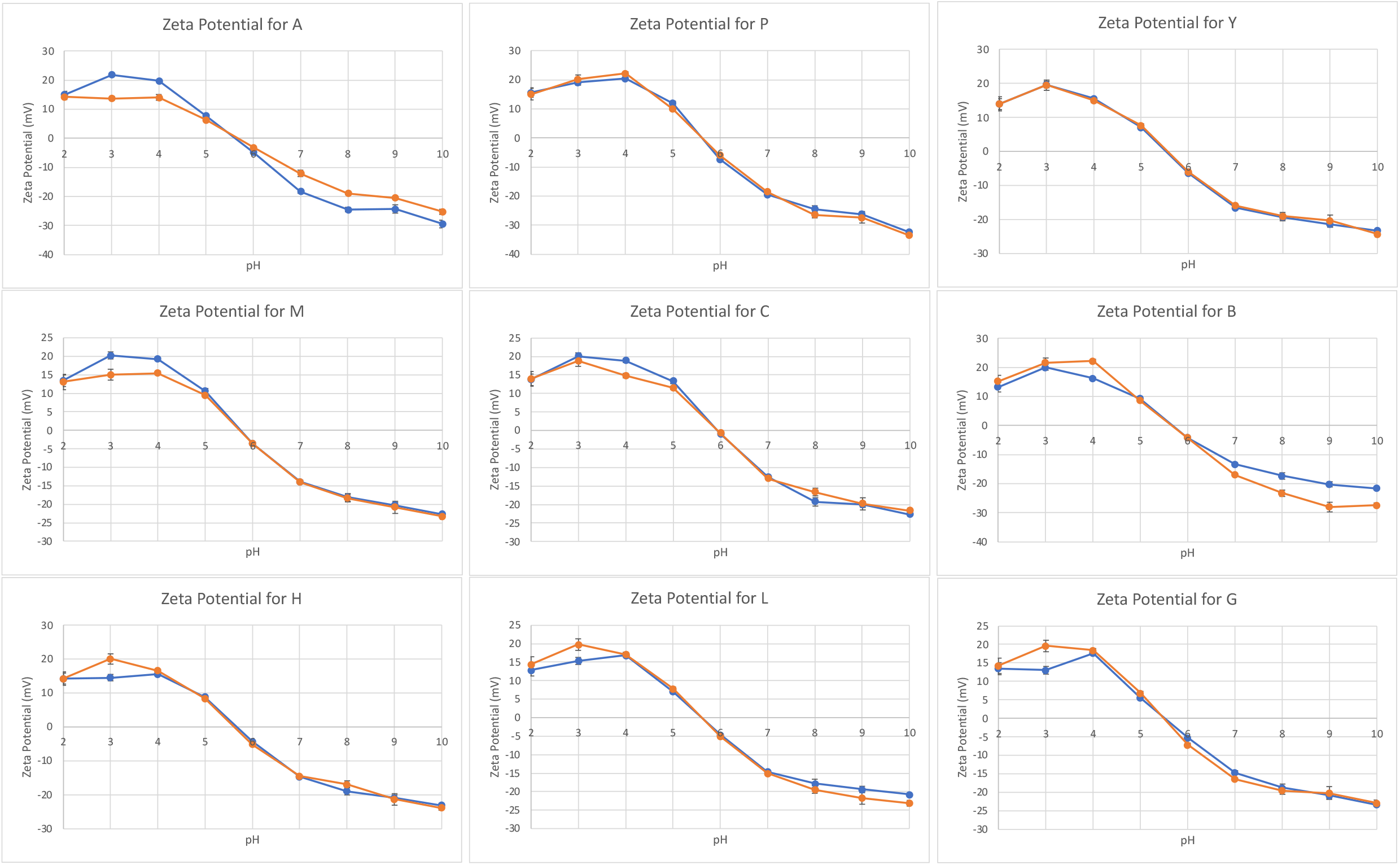
Zeta potential profiles of hemp seed protein isolates from each cultivar. Experiments were conducted in duplicate, and each measurement was conducted in triplicate

**Table S1.** Various cultivars of hemp seed underwent protein extraction to produce hemp seed protein isolates. The solubility of these protein isolates was tested at pH 3.0 – 7.0 and reported as percentages.

| Source | Protein solubility (%) |  |  |  |  |
| --- | --- | --- | --- | --- | --- |
|  | pH 3.0 | pH 4.0 | pH 5.0 | pH 6.0 | pH 7.0 |
| G | 23.42 ± 3.23 a | 15.20 ± 1.99 a | 15.77 ± 1.48 a | 17.49 ± 1.10 a | 17.50 ± 1.18 a |
| L | 19.79 ± 4.17 a | 11.92 ± 2.12 abc | 12.21 ± 1.97 abc | 14.26 ± 1.73 b | 14.50 ± 1.63 b |
| C | 26.47 ± 1.08 a | 13.96 ± 1.14 ab | 13.80 ± 1.09 ab | 13.97 ± 0.41 b | 12.66 ± 0.53 bc |
| M | 20.34 ± 2.72 a | 12.44 ± 1.04 abc | 12.60 ± 1.35 abc | 13.46 ± 0.49 b | 12.33 ± 0.82 bc |
| Y | 24.74 ± 2.05 a | 13.15 ± 1.46 abc | 12.92 ± 2.13 abc | 13.27 ± 0.80 b | 11.48 ± 1.15 cd |
| P | 25.72 ± 2.30 a | 10.98 ± 1.29 abc | 11.04 ± 1.38 bc | 11.61 ± 0.47 bc | 10.15 ± 0.96 cde |
| B | 25.53 ± 3.31 a | 12.13 ± 1.54 abc | 11.99 ± 1.62 abc | 12.09 ± 1.34 bc | 9.87 ± 0.95 cde |
| H | 22.22 ± 4.63 a | 9.57 ± 1.48 c | 9.13 ± 1.68 c | 10.39 ± 1.22 c | 9.13 ± 0.90 de |
| A | 21.92 ± 3.52 a | 10.21 ± 1.07 bc | 10.12 ± 0.99 bc | 9.92 ± 0.43 c | 7.65 ± 0.83 e |

Values denote the mean ± one standard deviation (n=3). Different letters in each column represent statistically significant differences within a given pH range as calculated by Tukey's HSD ( $p < 0.05$ ).
Abbreviations: G: GVA-H-20-1179; L: La Crème; C: Commercial hemp hearts; M: Magic Bullet 5; Y: Yuma Crossbow; P: Picolo; B: Bialobrzeskie; H: Hlukhivs'ki 51; A: Anka

